# A real-time PCR for the differentiation of typhoidal and non-typhoidal *Salmonella*

**DOI:** 10.1101/537654

**Authors:** Satheesh Nair, Vineet Patel, Tadgh Hickey, Clare Maguire, David R Greig, Winnie Lee, Gauri Godbole, Kathie Grant, Marie Anne Chattaway

## Abstract

Rapid and accurate differentiation of *Salmonella* spp. causing enteric fever from non-typhoidal *Salmonella* is essential for clinical management of cases, laboratory risk management and implementation of public health measures. Current methods used for confirmation of identification including biochemistry and serotyping as well as whole genome sequencing analyses, takes several days. Here we report the development and evaluation of a real-time PCR assay that can be performed directly on crude DNA extracts from bacterial colonies, for the rapid identification of typhoidal and non-typhoidal *Salmonella*.

This novel two-hour assay identifies the genus *Salmonella* by detecting the *ttr* gene, encoding tetrathionate reductase, and defines typhoidal *Salmonella* by the detection of *S*. Typhi and Paratyphi-specific gene combinations. PCR assay performance was determined using 211 clinical cultures of *Salmonella* (114 non-typhoidal and 97 Typhoidal strains) and 7 clinical *non-Salmonella* cultures. In addition, the specificity of the assay was evaluated *in silico* using a diverse in-house collection of 1882 *Salmonella* whole genome sequences. The real-time PCR results for 218 isolates and the genomic analysis of the 1882 isolates produced 100% sensitivity and 100% specificity (based on a 7 gene profile) for identifying typhoidal *Salmonella* compared to the *Salmonella* whole genome sequening identification methods currently used at Public Health England.

This paper describes a robust real-time PCR assay for the rapid, accurate identification of typhoidal and non-typhoidal *Salmonella* which will be invaluable for the urgent screening of isolates from symptomatic individuals, the safe processing of isolates in laboratories and for assisting the management of public health risks.

## Introduction

*Salmonella* are a diverse genus of gastrointestinal pathogens that cause a wide spectrum of disease from self-limiting gastroenteritis (non-typhoidal salmonellae, NTS) to systemic enteric fever (typhoidal salmonellae - *Salmonella enterica* serovar Typhi, Paratyphi A, B and C). Salmonellosis is global but typhoidal *Salmonella* are found mainly in sub-Saharan Africa and South Asia where enteric fever is endemic (1); although the detailed local surveillance data from endemic regions remains poor (2). A current concern is the increase in bacteraemia (and focal infections) associated with multi-drug resistant NTS infection in sub-Saharan Africa. In high income countries such as the UK, invasive NTS infection is mainly confined to immune-compromised hosts and so the major risks are local outbreaks of NTS through poor food hygiene and typhoidal infections associated with travel to endemic regions.

Diagnostic hospital microbiology laboratories make only a presumptive identification of *Salmonella* spp.: they do not usually hold a sufficient range of specific antisera for full identification and rapid identification systems, such as Matrix Assisted Laser Desorption/Ionisation-Time of Flight, Mass Spectroscopy, are unable to fully speciate *Salmonella*. In reference laboratories where definitive microbiological methods for the identification of *Salmonella* by serology and biochemistry (3) do exist, the turnaround times are often lengthy because of weak expression of the somatic (O), flagellar (H) and Vi polysaccharide surface antigens leading to incomplete or incorrect identification of the serovars. Whole Genome Sequencing (WGS) for *Salmonella* (4) has simplified the process for identifying *Salmonella* serovars substantially but still takes days rather than the hours. Currently there are no rapid diagnostic tests for informing clinical and public health management of enteric fever or for ensuring *Salmonella* isolates are processed appropriately with respect to laboratory safety.

In the UK, salmonellosis is a significant public health problem causing morbidity, financial loss due to sickness and absenteeism until clearance from infection for certain professions. The clinical management of salmonellosis patients depends on diagnosis. Enteric fever is treated with antibiotics but non-typhoidal *Salmonella* gastroenteritis is usually self-limiting. Invasive disease needs to be treated with antibiotics specific to the strain causing infection. In addition, the processing of isolates or specimens in the laboratory from patients with suspected diarrhoeal infection depends on the identification of the causal agent. In the UK microorganisms that pose a risk to human health are classified into one of four hazard groups based on their ability to infect healthy humans. The classification of these organisms allows the risk they pose to laboratory and healthcare workers to be controlled by implementing safety measures proscribed by law. *S*. Typhi, *S*. Paratyphi A, B and C are classified as a Hazard Group 3 (HG3) pathogens requiring processing in a specialised containment level 3 (CL3) laboratory (5). It is clear, therefore, that in order to treat patients effectively and protect healthcare and laboratory staff, the rapid identification of a patient as being infected with a typhoidal salmonella is critical.

At present there is no single rapid method to identify all typhoidal (HG3 in the UK) *Salmonella*, even though genomic data on the presence or absence of genes in both typhoidal and non-typhoidal *Salmonella* are in abundance. The *ttr* gene, encoding tetrathionate reductase has been used as a PCR gene target to detect and identify *Salmonella* since it is present in all *Salmonella* spp. (6). However, it is not intended to distinguish typhoid and non-typhoidal subspecies. A few potential candidate genes for identifying HG3 *Salmonella* have been described previously. For example, the *tviB* gene, encoding a Vi polysaccharide capsule, which is present in *S*. Typhi and *S*. Paratyphi C, (7) but not in *S*. Paratyphi A or *S*. Paratyphi B, can identify a subset of typhoidal *Salmonella* but doesn’t distinguish *S*. Typhi or *S*. Paratyphi C. (8). In order to differentiate *Salmonella* serovars causing enteric fever, additional genes are required. Nga *et al* (2010) proposed using *SPA2308*, encoding a hypothetical protein, for the detection of *S*. Paratyphi A and STY0201 (also known as the *staG* gene), encoding a putative fimbrial protein, for the detection of *S*. Typhi in clinical blood samples via PCR (9). Connor *et al* (2016) suggested that *S*. Paratyphi B (HG3) could be distinguished from *S*. Java (HG2) using two genes encoding Type III Secretion System (TTSS) effector proteins; *sseJ* and *srfJ* (10): with *S*. Paratyphi B possessing only *srfJ* but *S*. Java possessing both *sseJ* and *srfJ*. However, as *sseJ* is also absent in *S*. Typhi and *S*. Paratyphi A, this gene cannot be used to differentiate all HG3 serovars or used alone as an HG2 marker. A potential gene target for *S*. Paratyphi C identification is the *SPC0869* target, a gene encoding a hypothetical protein, shown to be present only in *S*. Paratyphi C (8). The use of this gene requires further assessment to ensure it is a unique target amongst the *S*. Paratyphi C population as only five serovars were investigated in the study by Lui *et al*., 2009.

The design of a PCR assay to identify *Salmonella* and differentiate HG2 and HG3 *Salmonella* requires a multi-targeted approach with defined gene profiles and rigorous validation. The aim of this study was to develop and validate a real-time PCR assay to distinguish HG3 (Typhoidal) and HG2 (Non-typhoidal) *Salmonella* and identify specific serovars of HG3 *Salmonella*.

## Methodology

### Bacterial strains

A total of 211 *Salmonella enterica subsp* I isolates, received at the Gastrointestinal Bacterial Reference Unit (GBRU), Public Health England (PHE) between 2008 - 2017, (Table 1a) were used in this PCR study. Representative HG2 isolates from the two most common serovars, *S*. Enteritidis and *S*. Typhimurium, as well as serovars that can be difficult to distinguish from HG3 isolates by traditional methods, including *S*. Dublin, *S*. Java and *S*. Choleraesuis, were selected (Table 1a). Assay specificity was further investigated by the inclusion of four *Shigella* isolates (*S. flexneri, *S*. sonnei, *S*. dysenteriae, *S*. boydii*) and three *E. coli* isolates (containing either *eae* or *stx* genes) as representatives to test the specificity against other *Enterobacteriaceae* that are occasionally misidentified by referring clinical laboratories using automated identification platforms (Supp table 1).

**Table 1:** Number and type of *Salmonella* serovars tested via molecular PCR and by GeneFinder. (a) Via PCR Footnote: *A random selection of HG2 ST containing sporadic gene targets were chosen. EPEC: Enteropaothogenic *E. coli*. STEC: Shiga toxin-producing *E. coli*. (b) Via GeneFinder

| Sequence Type (ST) | eBURST Group (EBG) | Serovar | Serotype | Hazard Group (HG) | No. |
| --- | --- | --- | --- | --- | --- |
| 1,2, 2173 | 13 | *Salmonella* Typhi | 9,12[Vi]:d: – | HG3 | 61 |
| 85, 129 | 11 | *Salmonella* Paratyphi A | 1,2,12:a:[1,5] | HG3 | 15 |
| 86 | 5 | *Salmonella* Paratyphi B | 1,4,[5],12:b:1,2 | HG3 | 15 |
| 146 | 20 | *Salmonella* Paratyphi C | 6,7,[Vi]:c:1,5 | HG3 | 6 |
| **Total HG3 *Salmonella*** |  |  |  |  | **97** |
| 11, 183 | 11, 183 | *Salmonella* Enteritidis | 1,9,12:g,m:– | HG2 | 14 |
| 19, 34, 36 | 19, 34, 36 | *Salmonella* Typhimurium | 1,4,[5],12:i:1,2 | HG2 | 14 |
| 10 | 10 | *Salmonella* Dublin | 1,9,12[Vi]:g,p:– | HG2 | 14 |
| 43, 88, 2545 | 43, 88, 0 | *Salmonella* Java | 1,4,[5],12:b:1,2 | HG2 | 13 |
| 2902, 3226, 139, 145 | 0, 0, 6,6 | *Salmonella* Choleraesuis | 6,7,:c:1,5 | HG2 | 6 |
| Variable* | Variable* | Selection of *Salmonella* ssp. from GeneFinder analysis* | Variable – see supplementary Table 1 | HG2 | 53 |
| **Total HG2 *Salmonella*** |  |  |  |  | **114** |
| 245, 152, 252, 7375 | CC245, 152, 145, 0 | *Shigella flexneri*, *S. sonnei*, *S. boydii*, *S. dysenteriae* | 3a, N/A, O6, O1 | HG2 | 4 |
| 11,29, 40 | CC11, 21, 40 | *E. coli* EPEC - eae +ve, STEC - stx1a, STEC - eae, stx2a | O55:H12, O77:H1, O157:H7 | HG2 | 3 |
| **Total of Non-*Salmonella*** |  |  |  |  | **7** |
| **Total of isolates tested** |  |  |  |  | **218** |

| Strains | No. |
| --- | --- |
| *Salmonella* Typhi | 556 |
| *Salmonella* Paratyphi A | 315 |
| *Salmonella* Paratyphi B | 53 |
| *Salmonella* Paratyphi C | 6 |
| HG2 Serovars | 952 |
| Non-Salmonellae | 7 |
| Total | 1889 |
| Strains | No. |
| No. of different Sequence Types | 480 |
| Subspecies I | 1821/1889 |
| Subspecies II | 14/1889 |
| Subspecies IIIa | 14/1889 |
| Subspecies IIIb | 29/1889 |
| Subspecies IV | 3/1889 |
| Subspecies V | 1/1889 |

### *Salmonella* whole genome sequence data

1882 *Salmonella* whole genome sequences (including the 211 *Salmonella* isolates), representing the diversity of Salmonellae tested by GBRU, were included in an *in silico* validation of the specificity of the selected target genes (Figure 1). This dataset included representative sequence types (ST) of the 19,221 strains validated and reported at GBRU between 2016-2017. The strains selected included all sub-species of *Salmonella* and the common (3 or more isolates received between 2016-2017 at PHE) *Salmonella* Serovars, representing in total 477 different sequence types (Table 1b, Supp table 1).

**Figure 1:**
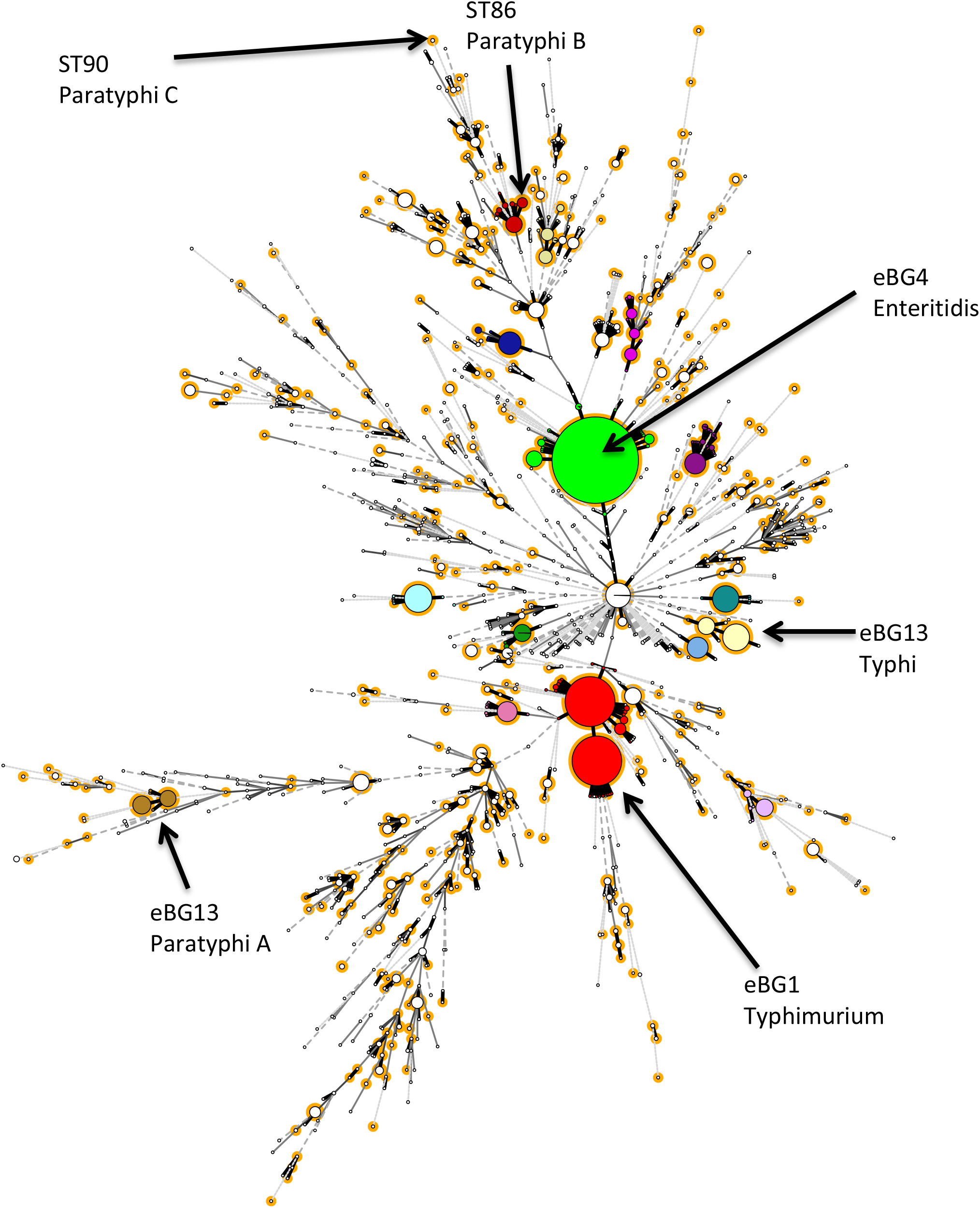
Selection of representative strains to test *in silico*. Footnote: Population structure of *Salmonella* received at PHE between 2016 – 2017 and strains tested for PCR in this study totalling to 19,221 strains. Colour coded by main eBURST groups (eBG), a representative strain (highlighted in orange) from each sequence types within an eBG containing 3 or more isolates were selected for *in silco* gene detection of the seven genes (*ttr, sseJ, srfJ, tviB, SPC0869, SPA2308 & staG*).

### DNA extraction and real-time PCR assays

DNA from 218 isolates was extracted via a crude extraction method in which a single colony from MacConkey agar [ThermoFisher Scientific, Waltham, USA] was inoculated into 490 μL of sterile distilled water in a screw cap microtube [Eppendorf, Hamburg, Germany] and placed in a boiling water bath for 20 minutes. Primers and probes for *ttr* (detection of all *Salmonella), tviB* (detection of *S*. Typhi and *S*. Paratyphi C), *SPA2308* (detection of *S*. Paratyphi A) and *staG* (detection of *S*. Typhi) were based on previous published studies (Table 2). Primers and probes for *SPC0869* (detection of *S*. Paratyphi C) *sseJ* and *srfJ* (detection of *S*. Paratyphi B) were designed using the PrimerQuest Tool V8 (https://www.idtdna.com/PrimerQuest/Home/Index) using sequences obtained from the NCBI nucleotide database (https://www.ncbi.nlm.nih.gov/nucleotide/) (Table 2).

**Table 2:**
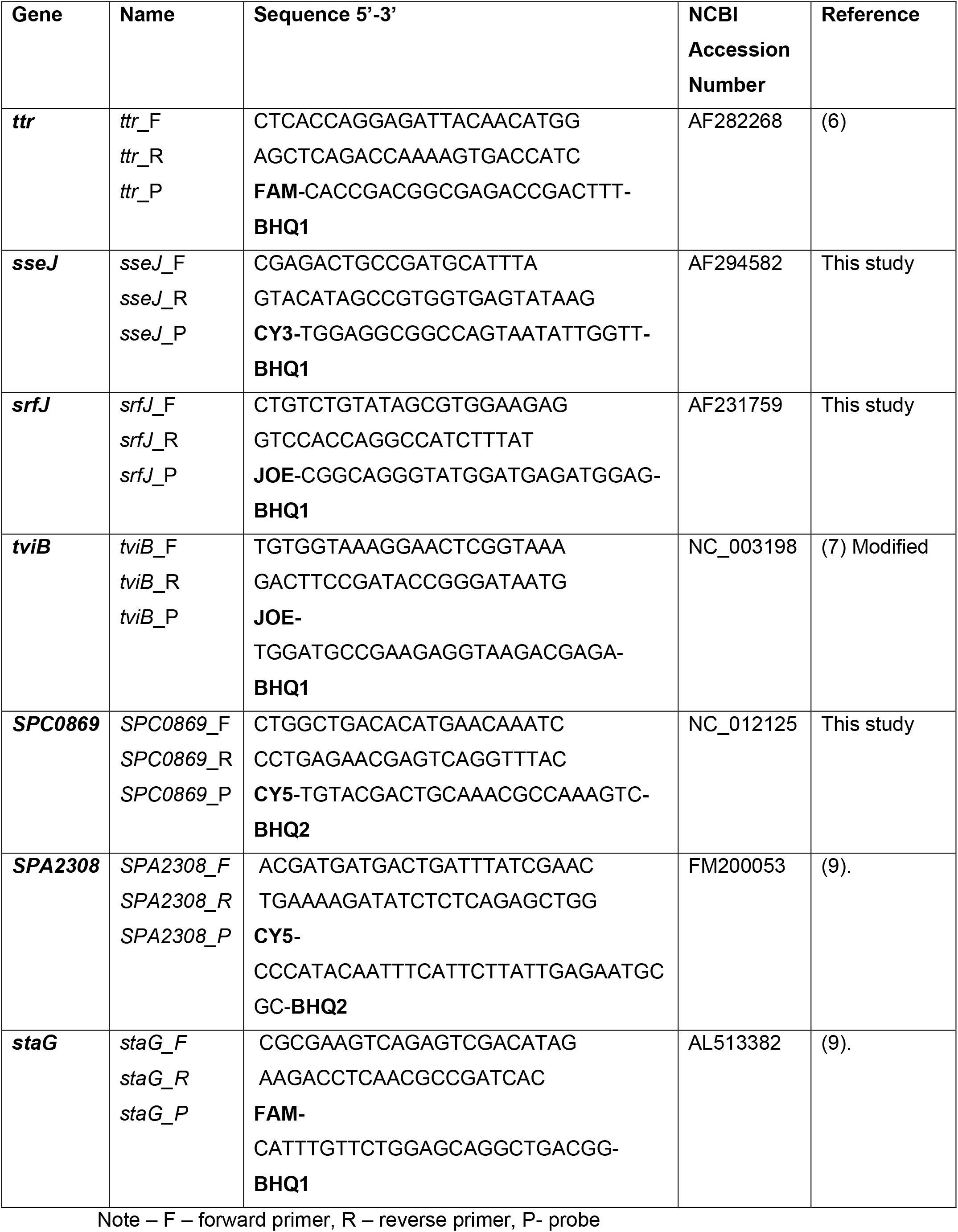
Primer and probe sequences used for each gene target with the fluorescent dye coloured (Colour of reporter related to spectrum of detection) and quenchers in bold (BHQ-black hole quencher). Footnote: Note – F – forward primer, R – reverse primer, P-probe.

The real-time PCR reported here was carried out as seven individual monoplex reactions but also worked as duplex and triplex PCR assays with interchangeable primers and probes targets (and probe dyes) depending on the target gene combination required. Mastermix for the monoplex assay consisted of 12.5 μL Takyon Low Rox probe mastermix [Eurogentec, Liège, Belgium], 8 μL Nuclease free water, 0.5 μL each of 20 μM forward and reverse primers, 1 μL of 5 μM probe and 2.5 μL DNA to a final reaction volume of 25 μL. A negative control was run with each PCR using 2.5 μL nuclease free water for the template [Severn Biotech, Kidderminster, UK] and the following positive controls were used: NCTC 8385 – *S*. Typhi (*ttr, tviB, staG*) NCTC 11803 - *S*. Paratyphi A (*ttr, SPA2308*), NCTC 8299 – *S*. Paratyphi B (*ttr, srfj*), NCTC 96 – *S*. Paratyphi C (*ttr, sseJ, tviB, SPC0869*), NCTC 6676 – *S*. Enteritidis (*ttr, sseJ*) and NCTC 14013 – *S*. Typhimurium (*ttr, sseJ, srfJ*). The PCR was run on the ABI Prism 7500 Real-Time PCR System [Applied Biosystems, Foster City, USA]. The conditions for the PCR were an initial activation of 95°C for 3 minutes, followed by 40 cycles of: Denaturation at 95°C for 30 seconds, Annealing at 60°C for 30 seconds, Extension at 72°C for 10 seconds. A positive result was assigned when a Ct value was achieved between 12-30 with a threshold set at 0.03ΔR.

Identification of HG3 *Salmonella* and differentiation from HG2 *Salmonella* was based on a profile of seven genes (Table 3). The molecular and/or *in silico* PCR identification was compared with the original identification of the serovar obtained via a combination of WGS identification, phenotype and serology carried out by the *Salmonella* laboratory as described previously (4) (Supp table 1).

**Table 3:** Gene profiles for the identification of *S*. Typhi and *S*. Paratyphi from other Serovars. Footnotes *#tviB* + means the strain is genotypically Vi positive. *Footnote: A proportion of HG2 Serovars will be positive for the *ttr* gene and a combination of targets that do not match any of the HG3 profiles (Supp table 1).

| Profile | *Salmonella* identification | *ttr* | *sseJ* | *tviB*# | *srfJ* | *SPC0869* | *SPA2308* | *staG* |
| --- | --- | --- | --- | --- | --- | --- | --- | --- |
| 1 | HG3 *Salmonella* Typhi | + | - | +/- | - | - | - | + |
| 2 | HG3 *Salmonella* Paratyphi A | + | - | - | - | - | + | - |
| 3 | HG3 *Salmonella* Paratyphi B | + | - | - | + | - | - | - |
| 4 | HG3 *Salmonella* Paratyphi C | + | + | + | - | + | - | - |
| 5 | HG2 Serovar* | + | + | - | +/- | - | - | - |
| 6 | Non-*Salmonella* spp. | - | - | - | - | - | - | - |

#*tviB* + means the strain is genotypically Vi positive

### PCR assay evaluation

The sensitivity and specificity of the *ttr, sseJ, srfJ, tviB, staG, SPA2308* and *SPC0869* primers and probes (Table 3) used in the real-time PCR assays were calculated according to Martin, 1984 (11).

In addition, PCR assay specificity was assessed by *in silico* genomic analysis using a diverse in-house WGS dataset covering the population structure of *Salmonella* (Figure 1). A total of 1882 *Salmonella* sequences (Supp table 1, Figure 1) which includes the 211 Salmonella isolates tested by PCR were screened for the presence of seven target genes *(ttr, sseJ, srfJ, tviB, staG, SPA2308* and *SPC0869)* using a PHE in-house bioinformatics tool called GeneFinder (developed by Doumith M, *et al*, unpublished). This tool takes paired-end Illumina FASTQ reads and aligns them to a reference sequence of the target genes, as a multi-FASTA file, (Accession numbers in table 2) using Bowtie2 v2.1.0 (12) and Samtools v1.0.18 (13) and determines metrics such as coverage, presence of indels (an insertion or deletion), amino acid alterations, presence of single nucleotide polymorphisms and overall sequence similarity of the test sequence to the reference gene sequence. Target genes were designated as present when sequences achieved a detection threshold of 80% sequence similarity to the reference gene, apart from *ttr* where the threshold was set at 70% sequence similarity, due to the size and variability of this particular gene. Any discrepant results between GeneFinder and the PCR were investigated further by assembling the sequence data using Spades v3.1.1 to default parameters and examining the variability of primer and probe binding sites.

Assay reproducibility was determined by testing 20 of the 211 *Salmonella* isolates in triplicate. Precision was evaluated by the standard deviation of Ct values of n=10 replicates of each of the positive controls for each target. Each target was assessed individually and as a multiplex in separate assay runs by different individuals and had the threshold set at 25% of the maximal fluorescence (ΔR) of each respective target.

## Results

### Comparison of real-time PCR and current PHE methods for distinguishing HG2 and HG2 Salmonellae

Of the 211 *Salmonella* isolates subjected to PCR identification, all gave the expected gene profile identification (Table 4, Supp table 1), matching the original identification, except for three *S*. Typhi isolates where the *tviB* gene was not detected. This was confirmed by *in silico* analysis (see below). Previous described ‘HG3’ gene targets *SPA0869, staG and SPA2308* were found sporadically in 41/114 (35%) of the HG2 *Salmonella* tested (two isolates had two HG3 gene targets present) confirming that use of single targets to differentiate HG3 from HG2 *Salmonella* is not appropriate (Table 5, Supp table 1). None of the 7 target genes were detected in the four *Shigellae* and three *E. coli* isolates that were tested.

**Table 4.**
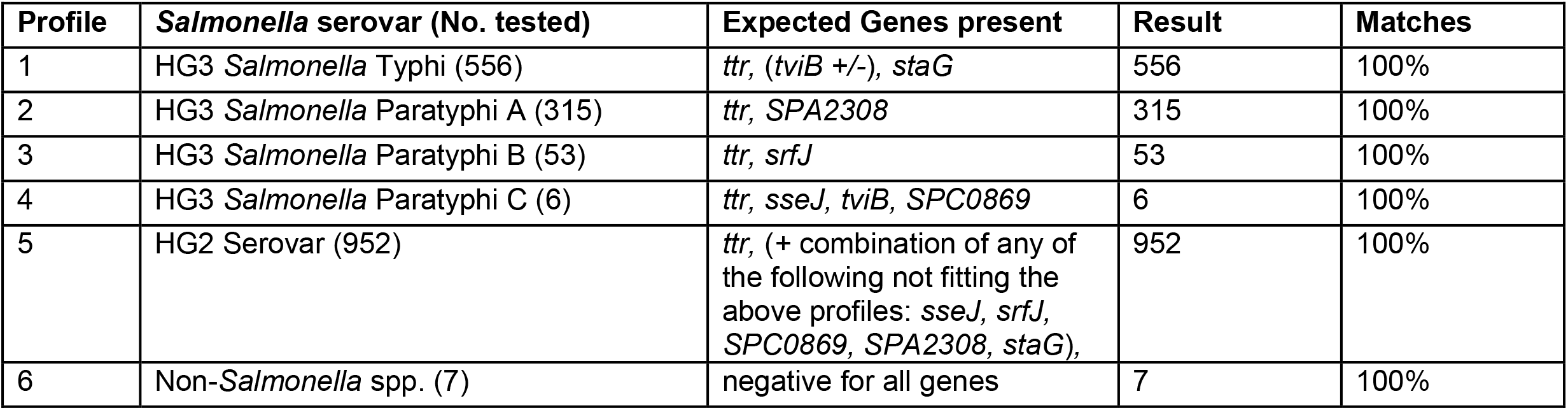
Summary of gene profile results.

**Table 5.**
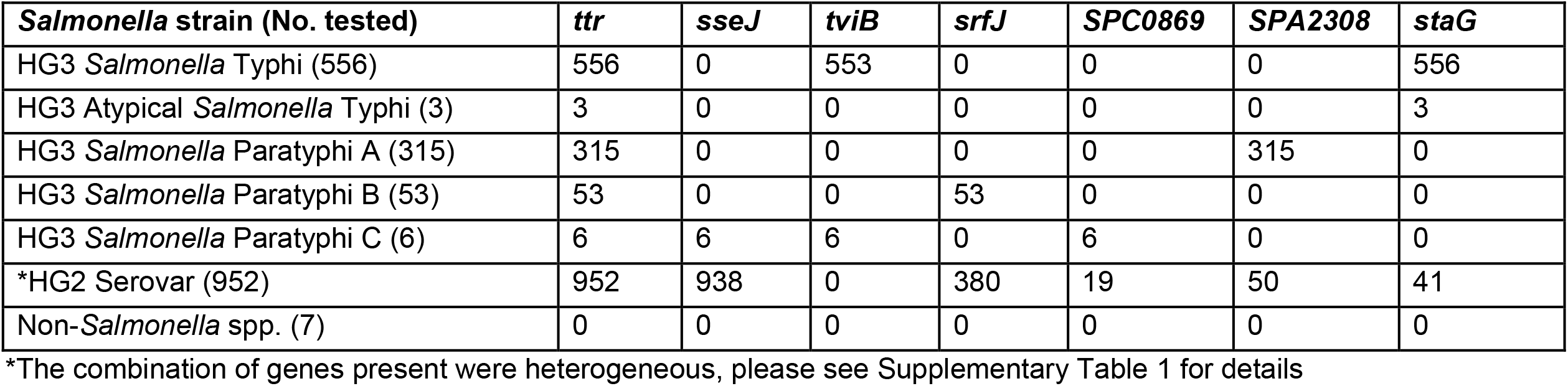
Summary of individual gene target results. Footnote *The combination of genes present were heterogeneous, please see Supplementary Table 1 for details.

### Whole genome sequencing *in silico* analysis

Of the 1882 Salmonella analysis subjected to *in silico* analysis, identification based on gene profiles (Table 3) matched the original identification but did highlight that individual gene targets could be found sporadically across the *Salmonella* population. *In silico* analysis identified 952/1882 non-typhoidal *Salmonella* isolates that were positive for *ttr* and a combination of other ‘HG3’ gene targets (Table 5, Supp table 1), designated as profile 5 (Table 4). None of the gene profiles of these isolates matched the designated HG3 profiles (profile 1-4) and thus our interpretation is that *ttr* positive strains with a profile not matching the HG3 profiles should be classified as HG2 *Salmonella* (Table 3, Table 4, Supp table 1).

As with the real-time PCR assay, the *Salmonella* processed via *in silico* analysis identified the three SPI-7 negative *S*. Typhi isolates. The real-time PCR and GeneFinder correctly identified the deletion of this gene.

In this study 8 of the 1882 sequences were positive by PCR and yet negative for the same gene by GeneFinder. Further *in silico* analysis revealed that the genes concerned had an intact primer and probe binding site, thus confirming the PCR result but variation outside of these regions resulted in average similarity values below the GeneFinder threshold value (Supp table 1).

### Reproducibility and precision of PCR assay

Reproducibility was assessed by performing the PCR 3 times on 20 isolates. Results indicated that the PCR was reproducible for differentiating between HG2 and HG3 salmonellae and for the identification of serovars within HG3 *Salmonella* (Supp table 1). The precision analysis demonstrated that five out of seven of the gene targets were considered precise (i.e. standard deviation <0.167). The following results show the gene, average Ct (and standard deviation): *ttr* - 25.12 (0.154), *sseJ* - 23.59 (0.127), *SrfJ* - 24.51 (0.179), *tviB* - 25.01 (0.115), *StaG* - 24.97 (0.121), *SPC0869* - 25.68 (0.142) and *SPA2308* - 20.59 (0.248). Both *SrfJ* and *SPA2308* have standard deviations above the 0.167 value that is considered precise. The explanation for this is that these two primer/probe sets are more susceptible to variation due to the *SrfJ* reverse primer having no G/C’s in the GC clamp therefore increasing the possibility of variable binding to the target gene. The *SPA2308* forward primer has less than 40% GC content making it more thermally variable and both reverse primer and probe’s do self-anneal and form hairpins. This is the case as the *SPA2308* gene has a very low GC content of 32.25% and as a result will lead to more variable results. Another important note is that this validation process occurred using boiled cells as the DNA extraction method (as this is the intended use for rapidity) and there is always the possibility of slight levels of PCR inhibition, in comparison to using purified DNA, which will also affect the precision results. The lower precision levels did not affect the molecular PCR in practice and was deemed suitable for use.

Reproducibility was not affected when targets were tested as a multiplex assay, however the precision assay in the molecular multiplex PCR proved to be better than the molecular monoplex reactions (Supp data 1).

### Sensitivity and specificity

Sensitivity and specificity were based on the 7 gene profiles (and not individual gene markers) detected by real-time PCR and GeneFinder (Table 4). It showed 100% sensitivity and specificity for the detection of HG3 *Salmonella* as compared to the routine reference identification by WGS and serotyping.

## Discussion

This study describes for the first time a robust real-time PCR assay for the specific identification of each of the four typhoidal *Salmonella* serovars: *S*. Typhi and *S*. Paratyphi A, B and C and is 100% reliable (Figure 1, Table 4, Table 5, Supp table 1). This assay was validated as a monoplex PCR providing the flexibility to use individual targets of interest but the assay was also found to work equally well as a multiplex assay (Supp data 1) and is now in use routinely at PHE. The rapid turnaround time of this PCR assay has potential for expediting the management of suspected cases of typhoid fever. With additional optimisation, the application of this assay could be extended to direct testing of clinical specimens (blood and stool) as well as food, water and environmental specimens. This would further increase the value of the assay although such use may risk the possibility of less isolates being referred to reference laboratories for further characterisation leading to loss of typing for surveillance purposes, including antimicrobial resistance monitoring, as well as outbreak detection and investigation. Thus, it is essential that isolates continue to be isolated and referred to reference laboratories.

Many assays for identifying typhoidal Salmonellae have been described previously but these are usually single gene methods with much lower specificity and sensitivity or are aimed at just one or two of the typhoidal serovars (9, 14–16). However, these important studies have provided input for the selection of candidate gene targets in designing a gene profile-based PCR assay, the validation of this PCR assay was strengthened by the use of WGS sequence data for high-through put testing on a more diverse collection of *Salmonella*.

*In silico* analysis has its limitations if relying on this approach as a sole method. Although PHE utilise a multilocus sequence type (MLST) based approach with genomic data for *Salmonella* identification (4), other organisations may use a gene based approach for *Salmonella* identification, the use of set thresholds with *in silico* testing in the current study on target genes (i.e. at what threshold is the test positive) may need to be flexible depending on the gene. Unlike detection via PCR, the entire target gene is evaluated using *in silico* analysis and therefore we can draw conclusions on the presence/absence of the target gene. However, selecting a threshold value (and therefore a percentage identify of a match) to which a gene is considered present or absent can be difficult. Discrepancies between, real-time PCR and genomic detection of target genes occur when a gene has less than the set threshold of sequence similarity. There were initially eight negative gene results using GeneFinder that were positive by PCR. These were due to a lower percentage of gene similarity and below the 80% set threshold (Supp table 1) and were positive for the presence of the gene (matching the PCR result). When mismatches between PCR and *in silico* methods occur, explicit consideration is required to ascertain if the PCR primer/probe binding region is intact and how much of the gene is present. Specifically, in our targets, *ttr* showed a large range of variability amongst isolates in terms of sequence similarity to the reference gene with five of eight of these samples having *ttr* <80% sequence similarity. After assessing the primer/probe binding sites of the genes, there were no discrepancies between GeneFinder and the PCR assay.

This current study showed that 17 of the 952 NTS isolates were only positive for a single gene target *(ttr* gene) (Supp table 1) and belonged to *Salmonella* subspp. III, IV and V. Therefore, most NTS *Salmonella* contain one or more of the other genes markers normally associated with typhoidal *Salmonella* (Table 5). This highlights that a single gene target method is not appropriate for distinguishing between typhoidal and non-typhoidal *Salmonella*, with a gene profile-based method being more accurate for identification and differentiation of typhoidal *Salmonella*. The reassuring finding, however, is that not one of the 935 NTS *Salmonella* had the same gene profile as the typhoidal (HG3) *Salmonella* profiles (Supp table 1, Table 3, Table 4).

Another notable observation is that three *S*. Typhi isolates from Pakistan lacked the 134kb SPI-7 pathogenicity island harbouring the *ViaB* operon *(tvi* genes – associated with the production of the Vi capsule). Although rare, absence of SPI-7 pathogenicity island including the *tvi* region in *S*. Typhi has previously been described (7). This is potentially an important public health finding as the current typhoid Vi polysaccharide vaccine stimulates immunity against the Vi capsule. It is known that SPI-7 negative (Vi-negative) *S*. Typhi can cause typhoid fever (17) and so there is a need to monitor the loss of the SPI-7 island in endemic regions where *S*. Typhi vaccination programs are being conducted (17). The assay described here could be used to monitor the emergence of Vi-negative *S*. Typhi through the emergence of *ttr* and *staG* positive *tviB* negative strains.

## Conclusion

In conclusion, this is the first real-time PCR assay that can rapidly distinguish between typhoidal ie *S*. Typhi, Paratyphi A, Paratyphi B and Paratyphi C (HG3) and non-typhoidal (HG2) *Salmonella* serovars. The assay has the ability to be implemented in diagnostic and reference laboratories globally as a safe and cost-effective way of differentiating *Salmonella*.

### Funding

This study was funded by PHE

## Acknowledgements

Thank you to Sarah Alexandra and Julie Russell from the National Collection of Type Cultures for providing positive control strains.

Thank you to Lailanie Aqunino from GBRU for support in undertaking PCR.

---

**Supplementary table 1** - Comparison analysis of reference methods versus *Salmonella* HG3 PCR and GeneFinder.

**Supplementary data 1** – Supplementary data on how this PCR was multiplexed into two triplexes and one monoplex. This data includes further precision data on the multiplex version of this PCR, the recipe used to make the mastermixes and associated tables and figures.

